# Diversity patterns across 1,800 chloroplast genomes of wild (*Oryza rufipogon* Griff.) and cultivated rice (*O. sativa* L.)

**DOI:** 10.1101/094482

**Authors:** Peter Civáň, Terence A. Brown

**Author notes:** corresponding author: Peter Civáň.

## Abstract

Cultivated Asian rice *(O. sativa* L.) comprises several groups with distinct ecological requirements and culinary uses. While the two subspecies of *O. sativa* – *indica* and *japonica* – have been subjected to a multitude of genetic and genomic analyses, less is known about the origins and diversity of the agronomically marginal groups – *aus* and aromatic rice. Here we reconstructed complete chloroplast genomes of over 1,800 accessions of wild and cultivated rice, including 240 *aus* and 73 aromatic varieties, and analysed the haplotype diversity of the taxonomic groups. We confirm the deep phylogenetic divergence between the main chloroplast haplotypes of *japonica* and *indica,* and reveal unique profiles of chloroplast diversity in *aus* and aromatic rice. Our results indicate that the latter two groups are not simple derivatives of *indica* and *japonica,* respectively, but originated from independent and/or reticulate domestication processes. Absence of phylogeographic patterns in the wild distribution of chloroplast haplogroups did not allow firm conclusions about geographic origins and the role of inter-group gene flow. Nonetheless, our results suggest that the domestication of *indica, japonica, aus* and aromatic rice operated on genetically different gene pools and followed different dynamics.

## Introduction

Cultivated Asian rice (*Oryza sativa* L.) is one of the oldest and most important staple crops worldwide. It is well established that *O. sativa* originated from a wild progenitor species *Oryza rufipogon* Griff., but the number of domestication events and the population genetic details of the process remain controversial (He et al. 2011; Yang et al. 2012; Molina et al. 2011; Huang et al. 2012; Civáň et al. 2015; Huang and Han 2015; Civáň and Brown 2016). Based on ecology, genetics and culinary properties, *O. sativa* can be divided into five groups – *japonica* (subdivided into tropical and temperate ecological groups), *indica, aus,* and aromatic rice (Garris et al 2005; Zhao et al. 2011; Civáň et al. 2015). *Japonica* varieties produce glutinous grain that is particularly sticky after cooking compared to long-grained *indica* and *aus* rice, the latter consisting of drought-tolerant, early-maturing varieties. Aromatic rice includes cultivars with specific flavors popular in Pakistan and northern India (basmati) and Iran (sadri). Reproductive separation of *O. sativa* and *O. rufipogon* is incomplete and gene flow between these species has been well-documented due to concerns of transgene escape from genetically modified rice (Song et al. 2003; Chen et al. 2004; Wang et al. 2006; Shivrain et al. 2007). An incomplete reproductive barrier was also described between *indica* and *japonica* rice (Harushima et al. 2002; Yang et al. 2012), but data on *aus* and aromatic crossability are missing.

The question of the genetic and geographic origins of the four rice groups *(indica, japonica, aus* and aromatic) is still debated. In recent years, the dominant view of the origins of *indica* and *japonica* has hypothesized that *japonica* was domesticated first in southern China and *indica* was derived later in other regions by hybridization of *japonica* with locally adapted wild rice (Huang et al. 2012). This domestication model is consistent with the notion that gene flow significantly shaped the genomes of *indica* and *japonica* (He et al. 2011; Yang et al. 2012), and recent analyses of rice genes apparently targeted by domestication are often interpreted accordingly (Hua et al. 2015; Oikawa et al. 2015; Si et al. 2016). Nonetheless, the hypothesis that some crucial domestication traits of *indica* have been derived from *japonica* was challenged by our group (Civáň et al. 2015). It also becomes apparent that the marginal *aus* crop has a unique domestication history (Zhao et al. 2011; Schatz et al. 2014; Civáň et al. 2015). Further genetic structure within the *aus* group was recently discovered by Travis et al. (2015) who differentiated two genetic subgroups within 250 *aus* cultivars, one of which is associated with the term ‘*boro*’, used to describe the winter growing season in Bangladesh and Assam. Although the aromatic varieties have been traditionally associated with the *japonica* group (Garris et al. 2005; Zhao et al. 2011), we suggested that aromatic rice was rather derived from crosses between *aus* and *japonica* (Civáň et al. 2015).

Besides the genetic diversity present in the nuclear genetic compartment, cytoplasmic (chloroplast and mitochondrial) genomes provide alternative sources of genealogical data often used for deep phylogenetic reconstructions (chloroplast genomes in plants), as well as phylogeography of populations (e.g. human mitochondrial genome). Cytoplasmic genomes are typically uniparentally inherited and are generally considered to be non-recombining units of genetic material, which means that their genealogical history is analogous to the genealogical history of a single gene. Consequently, the phylogeny reconstructed from chloroplast or mitochondrial genomes can differ from the species-or population-level phylogeny for reasons of lineage sorting, even in the total absence of introgression (Avise and Wollenberg 1997). Because of this, chloroplast genomes may not be reliable for species-tree reconstruction in recently diverged taxa, even when a sufficient number of phylogenetically informative characters are present. However, domestication is a special case of very recent and relatively rapid speciation typically accompanied by a severe bottleneck due to strong artificial selection. In such a case, the chloroplast genome diversity of a crop may be extremely reduced in respect to the diversity of the wild populations, and this reduction can provide indications about the course of domestication. Chloroplast uniformity across the entire crop can be interpreted as evidence of a single origin and a rapid domestication, while high chloroplast diversity indicates multiple origins or a lengthy domestication process with inter-group gene flow. Using this rationale, population-scale chloroplast datasets were utilized to infer single origins of domesticated sunflower (Wills and Burke 2006) and peanut (Grabiele et al. 2012), multiple domestications of lima bean (Andueza-Noh et al. 2013), as well as to make implications about intra and inter-specific hybridizations that contributed to the current genetic diversity of the domesticated species (Delplancke et al. 2011; Nikiforova et al. 2013; Li et al. 2013).

Chloroplast genome data has also been used to study the relationships among the *Oryza* species (Kim et al. 2015; Wambugu et al. 2015) and particularly the context of rice domestication (Kawakami et al. 2007; Tong et al. 2016; Kumagai et al. 2016). The datasets used either comprise short chloroplast DNA fragments reconstructed for hundreds of rice accessions (Kawakami et al. 2007; Kumagai et al. 2016), or involve the entire chloroplast genomes of dozens (Kim et al. 2015; Wambugu et al. 2015) to hundreds rice accessions (Tong et al. 2016). These studies revealed that *indica* and *japonica* rice carry generally distinct chloroplast lineages – each group being more closely related to particular wild haplotypes than to one another – supporting the hypothesis of multiple origins of *O. sativa* (Kawakami et al. 2007; Tong et al. 2016; Kumagai et al. 2016). However, *aus* and aromatic rice varieties are not represented in these datasets and the origin and diversity of the cytoplasmic lineages in these two groups remain unknown.

In this study, we utilized available whole-genome sequencing datasets (Huang et al. 2012; 3K RGP 2014) and reconstructed complete or nearly-complete chloroplast genomes of ∽1,800 *Oryza* accessions, including 240 *aus* and 73 aromatic accessions. We compared the chloroplast DNA diversity to the nuclear genome diversity and discuss the revealed patterns in the context of the domestication process and possible post-domestication gene flow.

## Results and Discussion

### *Chloroplast haplotypes of* Oryza *populations and inter-group admixture*

We used previously published Illumina whole-genome sequencing datasets produced by the shotgun approach (Huang et al. 2012; 3K RGP 2014) to perform a population-scale assembly and analysis of complete chloroplast genomes. We report here complete or near-complete chloroplast genomes for 1,823 accessions of wild (460 accessions) and cultivated rice (519 *indica,* 409 *temperate japonica,* 75 *tropical japonica,* 240 *aus/boro,* 73 aromatic; Table S1) assembled through a highly efficient yet simple bioinformatics pipeline. In the 1,823 assemblies, typically about 7.4% of the raw data mapped to the chloroplast genome, resulting in each base covered 601× and 189× on average for the *O. rufipogon* and *O. sativa* samples respectively (with 184.7 and 86.8 mean standard deviation of coverage; Table S1).

A 120,613-nucleotide long alignment of the reconstructed genomes revealed 216 polymorphic positions with >0.5% frequency in the entire super-population. Only a single position out of the 216 SNPs had three character states, indicating that the phylogenetic signal is not distorted by multiple substitutions and the dataset is generally free of substitutional saturation. The Ka/Ks ratio (0.24) calculated for all coding regions indicates purifying selection, although parts of some genes *(matK, rpoA, rpoC2)* may be evolving under positive selection. A data matrix of the 216 SNPs and 1,763 rice accessions containing 0.7% missing data was used to construct a median-joining network (Fig. 1). The 216 polymorphic positions define 107 distinct haplotypes that can be assigned into six major haplogroups (Fig. 1). In total, 67 haplotypes were found within wild rice and 55 haplotypes within cultivated rice.

**Fig. 1.**
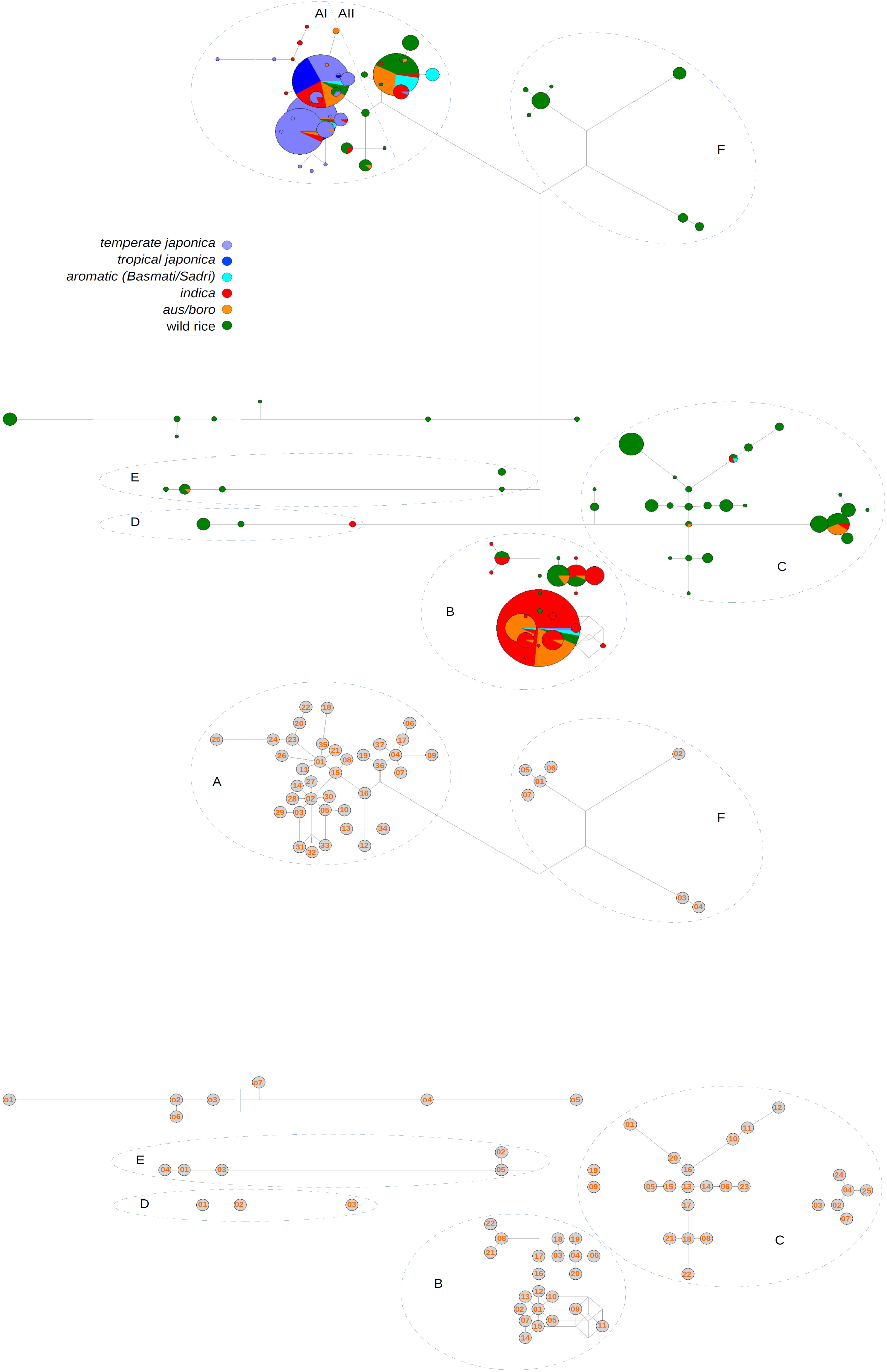
Median-joining network of rice chloroplast haplotypes. This is an unrooted network, however the root is most likely placed among the “o” (outgroup) haplotypes. This branch includes African *O. barthii* and *O. glaberrima* (o4), *O. longistaminata* (o7), *O. meridionalis* (o1-o3, o6) as well as *O. rufipogon* accessions sampled in Australia and Indonesia. Due to the uncertain root placement, the “o” haplotypes are not assigned to a formal haplogroup. (Upper panel) Six haplogroups designated A–F. Edge lengths are proportional to the number of polymorphisms separating the nodes and node sizes are proportional to the number of samples. As the group composition of haplogroup A is clearly heterogenous, we subdivided it further into haplogroup AI (typical for *japonica* accessions) and AII (*japonica* accessions absent). (Lower panel) Six haplogroups (A–F) and 107 haplotypes are labelled. Each node represents a distinct haplotype identified by the haplogroup letter and the node number. The haplotypes are numbered according to their ordered frequency, with the haplotype .1 being the most frequent. Edge lengths are proportional to the number of polymorphisms separating the nodes, except the edge between the haplotypes o3 and o7 (shortened).

Each cultivated group has a unique profile of chloroplast haplotypes. In temperate and tropical *japonica* we found 21 haplotypes in total, 20 of these in haplogroup A with three haplotypes being particularly common – A02 (31.4% of *japonica* samples), A03 (28.5%) and A01 (24.3%) – and 1 haplotype in haplogroup B (B01 – 1%). Within the *indica* group, we found 33 haplotypes in total, with haplotype B01 being the most common (60.9%). While 83.3% of *indica* samples fall within haplogroup B, 15.2% are found within haplogroup A, sharing several haplotypes with *japonica* (particularly the haplotype A01). The clear differentiation of the main *japonica* and *indica* haplogroups (haplogroups A and B, respectively) is in agreement with earlier studies that resolved the chloroplast genomes of these two groups in two different *O. rufipogon/sativa* clades (Kim et al. 2015; Wambugu et al. 2015). Despite the deep divergence of the *japonica* A-lineages and the *indica* B-lineages, a significant portion of *indica* accessions carries non-B haplotypes, and this observation is consistent with the findings of Kumagai et al. (2016).

Within the *aus* group, we found 18 haplotypes, among which B01, B02, A04 and A01 (34.2%, 23.3%, 16.3% and 10.8%, respectively) are the most frequent. Hence, the *aus* accessions carry the *indica*-like B haplotypes (B01 and its derivative B02), and the *japonica*-like A haplotypes (A01), but also haplotypes that are absent in *japonica* and *indica* (A04). Interestingly, the haplotype A04 is shared with aromatic rice, where 60.2% of accessions carry A04 or its derivative A09. Six additional haplotypes were found in the aromatic accessions, with A01, A02, A14 and B01 being relatively frequent (all approximately 10%).

Substitution rates for chloroplast genomes are relatively slow and have been estimated at 1.52×10^−9^ (Yamane et al. 2006) and 1.06×10^−9^ (Middleton et al. 2014) per site per year in grasses. Within the rice chloroplast network, the largest distance between wild haplotypes is 90 substitutions separating the nodes A03 and o1. The latter comprises *O. meridionalis* and *O. rufipogon* sampled from Australia and Indonesia, confirming the unique identity of Australian wild rice reported previously (Waters et al. 2011). The most frequent haplotypes of *japonica* and *indica* (A01 and B01, respectively) are separated by 40 substitutions, which represents 116,000–166,000 years of divergence when the substitution rates by Yamane et al. (2006) and Middleton et al. (2014) are used. This figure is lower than the *indica–japonica* divergence estimates obtained from nuclear loci (Vitte et al. 2004; Ma & Bennetzen 2004; Zhu et al. 2005), implying that either the rate of chloroplast evolution or the nuclear divergence time have been overestimated. On average, cultivated haplotypes are separated by only 0.27 substitutions from their closest wild haplotype, indicating that most of the observed diversity pre-dates domestication.

Among the cultivated groups, *aus* has the highest diversity, while *japonica* has relatively high haplotype diversity, but the lowest haplogroup diversity (Table 1). As expected, chloroplast diversity in *O. rufipogon* is higher than in any of the *O. sativa* groups, both in terms of haplotype and haplogroup diversity. The *O. rufipogon* haplogroups do not display a clear phylogeographic pattern, each haplogroup encompassing a large geographic area stretching from India to south-east Asia and China (Fig. 2). This observation is in agreement with Liu et al. (2015) and Kumagai et al. (2016), who found no correlation between genetic groups and geographic regions. Since the divergence of the haplogroups precedes the last glacial maximum and perhaps also the previous interglacial (implied by the number of substitutions and the rate of chloroplast evolution), the absence of any phylogeographic pattern is probably due to repeated extinctions and recolonizations of wild populations during Quaternary glaciation cycles. Unfortunately, the wide distribution of the chloroplast haplogroups in the wild does not allow any inference of the geographic origins of the cultivated groups.

**Table 1.**
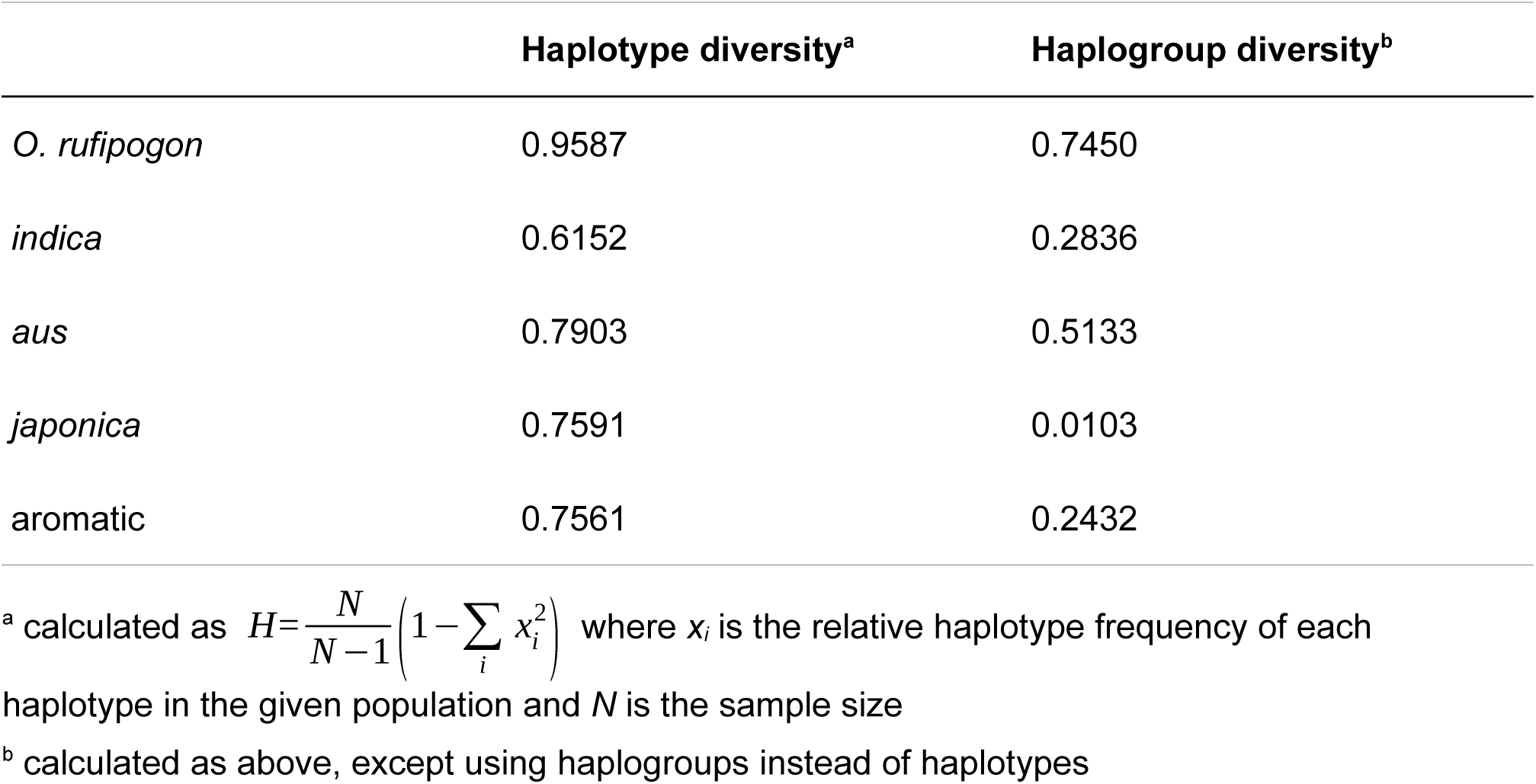
Chloroplast diversity in wild and cultivated rice.

|  | Haplotype diversity<sup>a</sup> | Haplogroup diversity<sup>b</sup> |
| --- | --- | --- |
| *O. rufipogon* | 0.9587 | 0.7450 |
| *indica* | 0.6152 | 0.2836 |
| *aus* | 0.7903 | 0.5133 |
| *japonica* | 0.7591 | 0.0103 |
| aromatic | 0.7561 | 0.2432 |

<sup>a</sup> calculated as $H = \frac{N}{N-1} \left( 1 - \sum_i x_i^2 \right)$ where $x_i$ is the relative haplotype frequency of each haplotype in the given population and $N$ is the sample size
<sup>b</sup> calculated as above, except using haplogroups instead of haplotypes

**Fig. 2.**
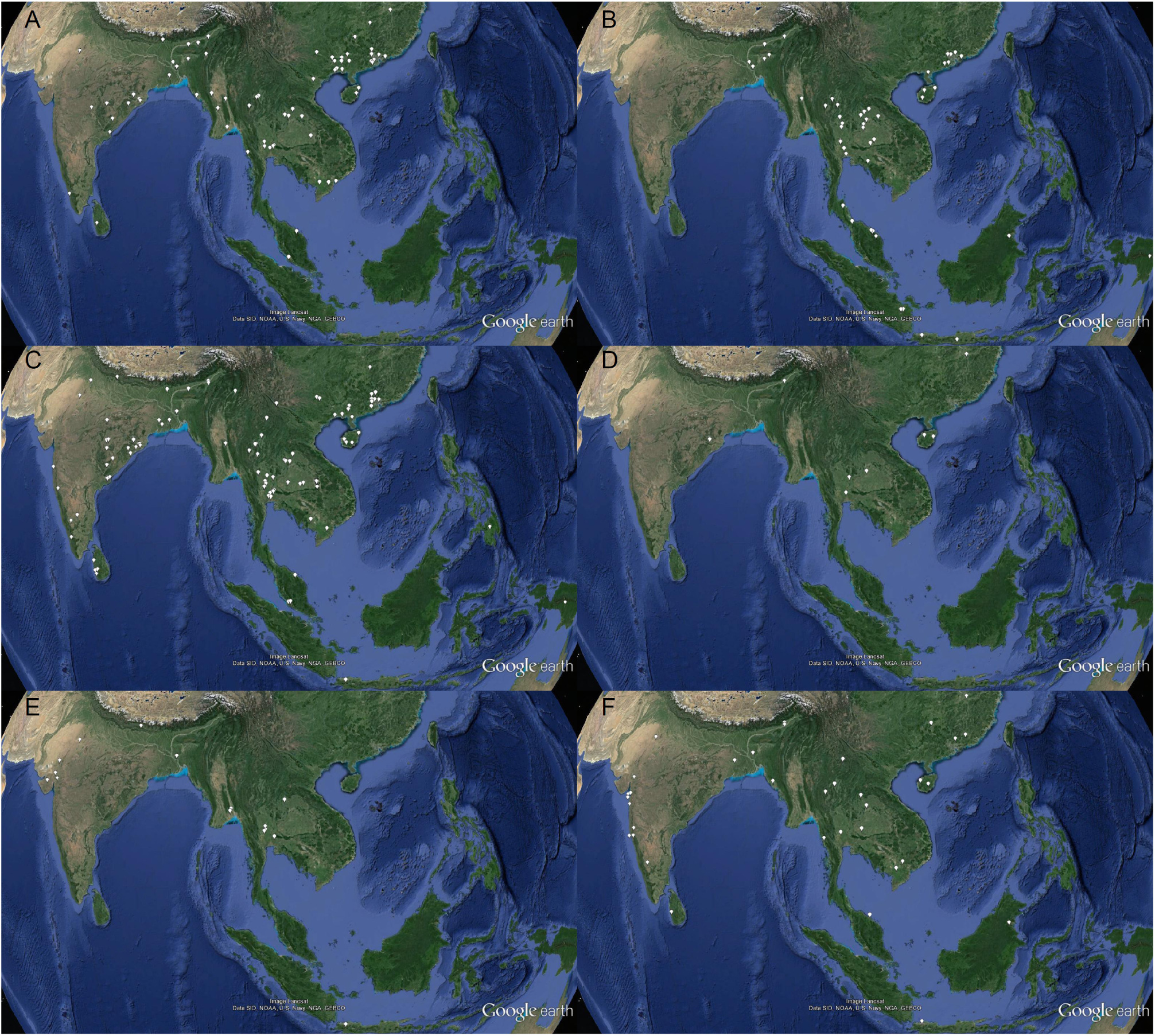
Geographic distribution of *O. rufipogon* chloroplast haplogroups. (A–F) Haplogroups A–F. A single map location may represent multiple samples. One Indian and 17 Chinese accessions are not shown due to their geographic data being unavailable. Maps prepared in Google Earth v7.1.5.1557.

A principal component analysis (PCA) performed on a genome-wide collection of nuclear single nucleotide polymorphisms (SnPs) revealed differences between the nuclear and chloroplast diversity patterns. The four cultivated groups are well differentiated and form clusters associated with three groups of *O. rufipogon* (Fig. 3A). All cultivated groups are differentiated by the first two eigenvectors with statistical significance, except for *aus* and *indica* along the first eigenvector. However, when the rice accessions in the same analysis are recoded according to their chloroplast haplogroups, the PCA clusters become heterogenous, indicating that the chloroplast haplogroups poorly represent the diversity of the nucleus. This is probably a reflection of the non-recombining nature of the chloroplast genome that passes through generations as a single locus strongly affected by lineage sorting (Avise and Wollenberg 1997). Discrepancies between the nuclear and cytoplasmic phylogenetic signals were also observed in other crop-wild systems (Nikiforova et al. 2013; Delplancke et al. 2013), which warns against phylogenetic inference solely based on chloroplast data from recently diverged species or populations.

**Fig. 3.**
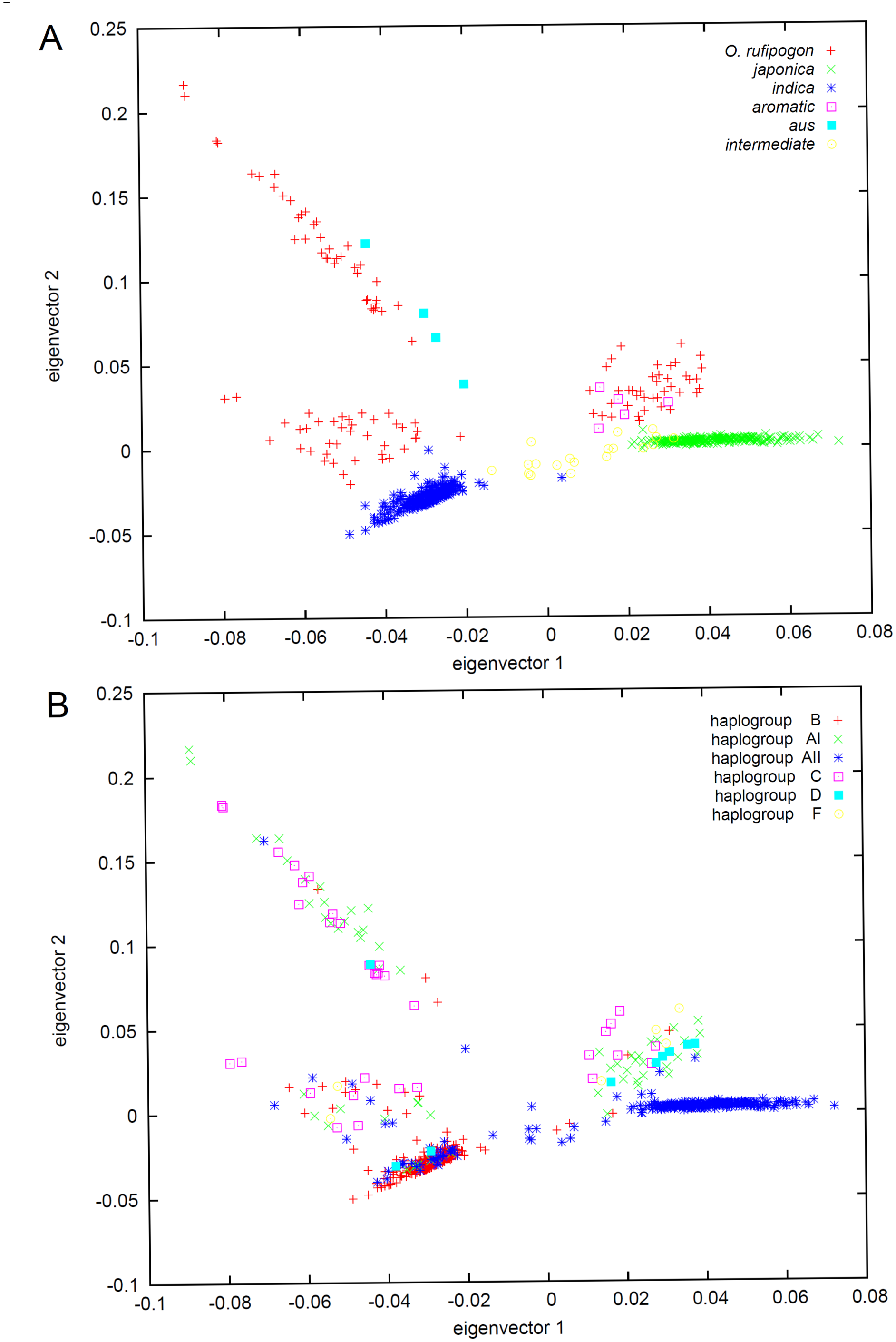
PCA of the cultivated groups and the identified ancestral gene pools. A total of 5,759,208 genome-wide nuclear SNPs were transformed into 700 axes of variation. The first two axes represent 9.7% of the total variation. (A) The first two eigenvectors with rice accessions coloured according to their domestication class. (B) The first two eigenvectors with rice accessions coloured according to their chloroplast haplogroups. It is apparent that rice samples with unrelated chloroplast haplogroups can be very similar at the nuclear level (note the dense clustering of *indica* samples carrying different haplotypes).

Nonetheless, the network of chloroplast diversity (Fig. 1) reveals one general distinction among the cultivated groups – while practically all *japonica* accessions have closely related haplotypes (98.8% derived from a single node A15), the other three groups contain rather unrelated haplotypes in non-negligible proportions. This observation is consistent with previous findings of generally lower levels of genetic diversity in the nuclear genome of *japonica* compared to *indica* (Caicedo et al. 2007; He et al. 2011; Huang et al. 2012) and suggests differences in the population dynamics during or after domestication. Two general models can be hypothesized to explain the observed pattern. (i) The chloroplast haplotypes of the cultivated groups were inherited vertically from multiple progenitor populations that differed in their genetic diversity. Under this model, the genetic basis of *japonica* was extremely narrow – either due to a very small ancestral population or a bottleneck caused by strong artificial selection – while the ancestral gene pools of *aus, indica* and aromatic rice were genetically wider. Under the second model (ii), the (chloroplast) diversity of *indica, aus* and aromatic rice was enriched by post-domestication admixture. Postdomestication admixture can, however, operate under two different scenarios – the cultivated groups were either domesticated independently from defined wild populations and subsequently admixed with other cultivated/wild gene pools, or all cultivated groups are derived from a single domestication event followed by very dynamic wild admixture events leading to the formation of distinct cultivated groups. The latter scenario was proposed by Huang et al. (2012) who suggest that *japonica* rice was the originally domesticated crop from which the other *O. sativa* groups were derived. Interestingly, the *japonica*-typical haplotype A1 is found in non-negligible frequencies in all cultivated groups, which allows the speculation that *japonica* played some role in the ancestry of *indica, aus* and aromatic rice. However, since the A haplogroup (and more specifically the AI haplogroup) is widely distributed in wild rice populations (Fig. 2A, Fig. 3), it is equally possible that all cultivated groups obtained A-haplotypes vertically from their wild progenitors. The haplotype profiles observed here can also be interpreted in accordance with the alternative domestication scheme we proposed recently (Civáň et al. 2015), in which the *indica, japonica* and *aus* groups were domesticated independently, *indica* and *aus* having wider and partially overlapping ancestral gene pools, while aromatic varieties arose from hybridizations between *japonica* and *aus*. Independent origins of *indica* and *japonica* are documented here by the deep divergence of their prevalent haplotypes, while the close association of *aus* and aromatic rice is supported by sharing the A04 haplotype that is frequent and almost exclusive in these two cultivated groups. The scenario of three independent origins of *O. sativa* and the hybrid ancestry of the aromatic varieties is also supported by the PCA of nuclear variation, where three clusters of *O. rufipogon* are associated with different cultivated groups and the aromatic varieties are positioned in between the *japonica* and *aus* clusters (Fig. 3A).

In summary, we provide a comprehensive picture of chloroplast diversity in *O. rufipogon* and *O. sativa* populations. The possibly strong effect of lineage sorting on the genealogical patterns of chloroplast genomes coupled with the absence of a phylogeographic pattern in wild rice does not permit definite conclusions about geographic origins of the cultivated groups or the exact role of inter-group gene flow. Nonetheless, our results provide further support to the notion that *japonica* has a more limited genetic basis than the other cultivated groups. Furthermore, the unique profiles of chloroplast haplotypes observed in *aus* and aromatic rice indicate that these groups are not simple derivatives of *indica* and *japonica,* but rather originated from independent and/or reticulate domestication processes.

## Materials and Methods

### Reconstruction and analysis of chloroplast genomes

Raw sequencing data for 1,823 wild and cultivated rice accessions (Table S1) published by Huang et al. (2012) and the 3k RGP (2014) were downloaded from the Sequence Read Archive (http://ncbi.nlm.nih.gov/Traces/sra/) and converted into FASTQ format using the fastq-dump command in sratoolkit 2.3.5. The full FASTQ dataset consisted of 12.0 billion Illumina reads totalling 3.1 TB. For each of the 1,823 accessions, reads significantly matching known rice chloroplast genomes (E-value <1e-5) were extracted using the interleave_pairs and filter_by_blast scripts from the seq_crumbs-0.1.9 package. Adapter contamination and low-quality regions were removed from the matching reads by Trimmomatic-0.33 (Bolger et al. 2014). These procedures yielded 721 million chloroplast-matching reads, totalling 256 GB, equating to >10-fold reduction of the original data. The filtered and trimmed datasets were imported into Geneious 6.1 (Biomatters; http://geneious.com) and individually mapped onto the Nipponbare chloroplast genome (KM088016) used as a reference (5 mapping iterations; maximum of 5% mismatches and 10% gaps per read; maximum gap size set to 100; index word length 13; only paired reads matching within the expected distance were mapped). The reconstructed chloroplast genomes with total quality scores are deposited at DOI: 10.5281/zenodo.61977. The second copy of the inverted repeat region was removed from the reconstructed sequences, which were then aligned using MAFFT v7 (Katoh et al. 2002) implemented in Geneious. Chloroplast genomes that were difficult to align due to a high proportion of missing data were further removed and the entire alignment manually corrected. The Ka/Ks ratio was calculated for a sample of 1,352 accessions with no missing data or ambiguities in their coding sequences using DnaSP (Librado and Rozas 2009). Polymorphic sites with frequencies <0.005 in the entire superpopulation were extracted from the alignment, treating all gaps as missing data and ignoring inversions and di-/poly-nucleotide changes. The resulting data matrix consisted of 216 sites and 1,763 rice accessions (461 accessions from wild *Oryza* species; 507 *indica;* 407 *temperate japonica;* 240 *aus;* 75 *tropical japonica;* 73 aromatic), with 0.7% missing data points in total. The data matrix was manually converted from FASTA into RDF format and imported into Network 4 (Bandelt et al. 1999) where a median-joining network was constructed. The accessions with intermediate phenotypes were not included in the network, but their haplogroups were recorded and are listed in Table S1. Chloroplast divergence times were calculated using the equation *T*=*D_XY_*/2*I*, where *T* is the divergence time, *D_XY_* is the proportion of differences observed between the sequences *X* and *Y*, and *l* is the rate of nucleotide substitution per site per year (Nei, 1987).

### PCA of nuclear variation

The complete genotype dataset for 1,529 wild and cultivated rice accessions consisting of ∽8 million SNPs from all twelve rice chromosomes published by Huang et al. (2012) was downloaded from the Rice Haplotype Map Project database (http://www.ncgr.ac.cn/RiceHap3). Individuals with >75% missing data points were removed from the dataset and the accessions of wild rice were further reduced to contain only 120 genotypes that represent the progenitor gene pools of *indica, japonica* and *aus* (Civáň P, unpublished data). Characters showing no variation in this subset of samples were removed, yielding a SNP matrix with 701 individuals and 5,759,208 positions. The PCA was performed with smartpca (Patterson et al. 2006), without outlier exclusion and inferring genetic distance from physical distance.

## Acknowledgements

This work was supported by European Research Council grant 339941 awarded to TAB.

